# Pharmacological depletion of microglia protects against alcohol-induced corticolimbic neurodegeneration during intoxication in male rats

**DOI:** 10.1101/2024.11.04.621917

**Authors:** Erika R. Carlson, Jennifer K. Melbourne, Kimberly Nixon

## Abstract

Excessive alcohol use damages the brain, especially corticolimbic regions such as the hippocampus and rhinal cortices, leading to learning and memory problems. While neuroimmune reactivity is hypothesized to underly alcohol-induced damage, direct evidence of the causative role of microglia, brain-resident immune cells, in this process is lacking. Here, we depleted microglia using PLX5622 (PLX), a CSF1R inhibitor commonly used in mice, but rarely in rats, and assessed cell death following binge-like alcohol exposure in male rats. Eleven days of PLX treatment depleted microglia >90%. Further, PLX treatment prevented alcohol-induced neuronal death in the hippocampus and rhinal cortices, as the number of FluoroJade-B-positive cells (dying neurons) was reduced to control diet levels. This study provides direct evidence that alcohol-induced microglial reactivity is neurotoxic in male rats. Improved understanding of alcohol-microglia interactions is essential for developing therapeutics that suppress pro-cytotoxic and/or amplify protective microglia activity to relieve alcohol-related damage.

## Introduction

Excessive alcohol consumption, characteristic of an alcohol use disorder (AUD), causes neurodegeneration as observed in MRIs of men with AUD (reviewed in Zahr and Pfefferbaum, 2017) and neuronal death in rodent models of alcohol exposure (Kelso et al., 2011; see Crews and Vetreno, 2014 for review). Microglia, immuno-competent brain cells with varied roles in neurodegenerative disease, are implicated as the primary mediators of the neuroimmune response to alcohol (Qin and Crews, 2012; Erickson et al., 2019; Melbourne et al., 2019). Postmortem brains from individuals with an AUD show increased microglia/macrophage markers such as ionized calcium-binding adapter molecule 1 (Iba1) and monocyte chemoattractant protein-1 (He and Crews, 2008) that correspond to similar findings in rodent models (Qin and Crews, 2012; Marshall et al., 2013; Peng et al., 2017; reviewed in Melbourne et al., 2019). The direct effects of alcohol on microglial cytokine gene expression have been shown *in vitro* (mouse BV2 cells; Walter and Crews, 2017), which were blunted by depletion of microglia in rat organotypic slice cultures (Coleman et al., 2020). However, there is no direct evidence of the causal role of microglia in alcohol-induced neurotoxicity. Therefore, we examined the role of microglia on alcohol-induced neurodegeneration in a rat model of an AUD by depleting microglia with PLX5622, a highly selective antagonist of colony-stimulating factor-1 receptor (CSF1R), a receptor kinase critical for survival and proliferation of microglia/macrophages (Riquier and Sollars, 2020; see Green et al., 2020).

## Methods

### Animal model

Male Sprague-Dawley rats (*n* = 34; 336.5 ± 3.6g; Charles River Laboratories, Raleigh, NC, USA) were group housed on a 12 h light/dark cycle with *ad libitum* food and water, except during diet administration. One ethanol-vehicle rat was excluded due to excessive FJB+ staining (outlier). To deplete microglia, rats received i.p. injections of CSF1R antagonist PLX5622 (PLX; MedChemExpress, Monmouth Junction, NJ, USA; 50 mg/kg) suspended in vehicle (5% DMSO, 20% Kolliphor RH40, and saline; 10 mL/kg; Riquier and Sollars, 2020) every 12 h for 11 days. In pilots, PLX-integrated chow was unpalatable and thus insufficient for microglia depletion. During the last 4 days of PLX treatment, rats received ethanol intragastrically every 8 h for 4 days in a Majchrowicz paradigm (Kelso et al., 2011; Marshall et al., 2013): following a 5 g/kg priming dose of ethanol (25% w/v in Vanilla Ensure Plus^®^, Abbott Laboratories, Abbot Park, IL, USA) or isocaloric (dextrose) control diet, doses were titrated (0-5 g/kg) based on intoxication behavior (i.e., a more intoxicated rat receives less ethanol). Blood ethanol concentrations (BEC; mg/dL) were determined by AM1 Alcohol Analyzer (Analox Instruments, Stourbridge, UK) from tail blood collected at “peak” intoxication (90 min post-7^th^ dose). Following the last dose (3-6 h post), rats were overdosed with anesthetic (Fatal-plus^®^, Vortech Pharmaceuticals, Dearborn, MI, USA), perfused transcardially with phosphate-buffered saline (PBS), and brains were extracted and one hemisphere post-fixed overnight in 4% paraformaldehyde at 4°C.

### Histology

Twelve series of 40 μm sections were obtained on a vibrating microtome (Leica VT1000S, Wetzlar, Germany) and stored in cryoprotectant at -20°C. Subjects were coded so that experimenters were blind to treatment. Free-floating, diaminobenzidine (DAB) immunohistochemistry for Iba1, a calcium binding protein expressed in all microglia/macrophages [1:1000 rabbit anti-Iba1 (FUJIFILM Wako, Osaka, Japan; RRID:AB_839504) blocked in 3% goat serum], was performed on one series of sections using Vector Laboratories (Newark, CA, USA) reagents as reported previously (Marshall et al., 2013): biotinylated goat anti-rabbit secondary antibody, Vectastain Elite ABC kit, and DAB Substrate kit. An adjacent series of sections was mounted to Superfrost Plus^®^ slides and stained for neurodegeneration with FluoroJade-B (FJB; Histo-Chem Inc., Jefferson, AR, USA) as reported (Kelso et al., 2011). All slides were coverslipped with Cytoseal^®^ (Thermo-Fisher Scientific, Waltham, MA, United States).

### Data Acquisition and Analysis

For Iba1, hippocampus (Bregma –2.30 to –4.50 mm; 3-5 sections/brain) and rhinal cortices (Bregma –3.00 to –7.68 mm; 4-8 sections/brain) were imaged on an Olympus BX43 microscope with DP73 camera and cellSens v3.1 (Olympus, Center Valley, PA, USA). The number of Iba1+ cells was estimated in ImageJ (macro available upon request) and reported as mean cells per field of view (FOV) ± standard error of the mean (SEM). FJB+ cells were counted in the hippocampus and rhinal cortices (Bregma –3.00 to –7.68 mm) on an Olympus BX51 microscope with epifluorescence (488λ cube for blue light excitation) and images acquired via DP70 camera (Olympus) and Stereo Investigator v2021.1.3 (MBF Bioscience, Williston, VT, USA). FJB+ cells are presented as mean cells per section ± SEM.

Data analysis was conducted in RStudio v2023.12.1.402 (Posit team, Boston, MA, USA) with R v4.2.2 (R Core Team, Vienna, Austria) using rstatix, afex, and emmeans packages (script available at https://github.com/ecarlson5683/carlson-2024-JNP-plx-ethanol) with visualization in Prism v10.0.3 (GraphPad, La Jolla, CA, USA). Intoxication behavior, ethanol dose, and BECs were analyzed by Welch’s t-test. Cell counts were analyzed by two-way ANOVA: factors of drug (PLX *vs* vehicle) and diet (ethanol *vs* control). Posthoc pairwise comparisons with Bonferroni correction were performed when appropriate. Significance was accepted at *p* < 0.05. Iba1+ cell count percent loss was calculated as 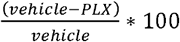.

### Results

PLX5622 successfully depleted microglia >90% in the hippocampus and rhinal cortex (Figure 1). In the hippocampus, two-way ANOVA revealed a main effect of drug (F_1,29_ = 164.1, *p* < 0.0001) on Iba1+ cells. Posthoc comparisons confirmed fewer Iba1+ cells with PLX versus vehicle treatment in both ethanol (92% depletion; *p* < 0.0001) and control (89% depletion; p < 0.0001) diet groups. Similarly, in the rhinal cortex, there was a main effect of drug (F_1,29_ = 262.1, *p* < 0.0001), with fewer Iba1+ cells in PLX-treated rats versus vehicle in both ethanol (95% depletion; *p* < 0.0001) and control diet groups (92% depletion; *p* < 0.0001).

**Fig. 1.**
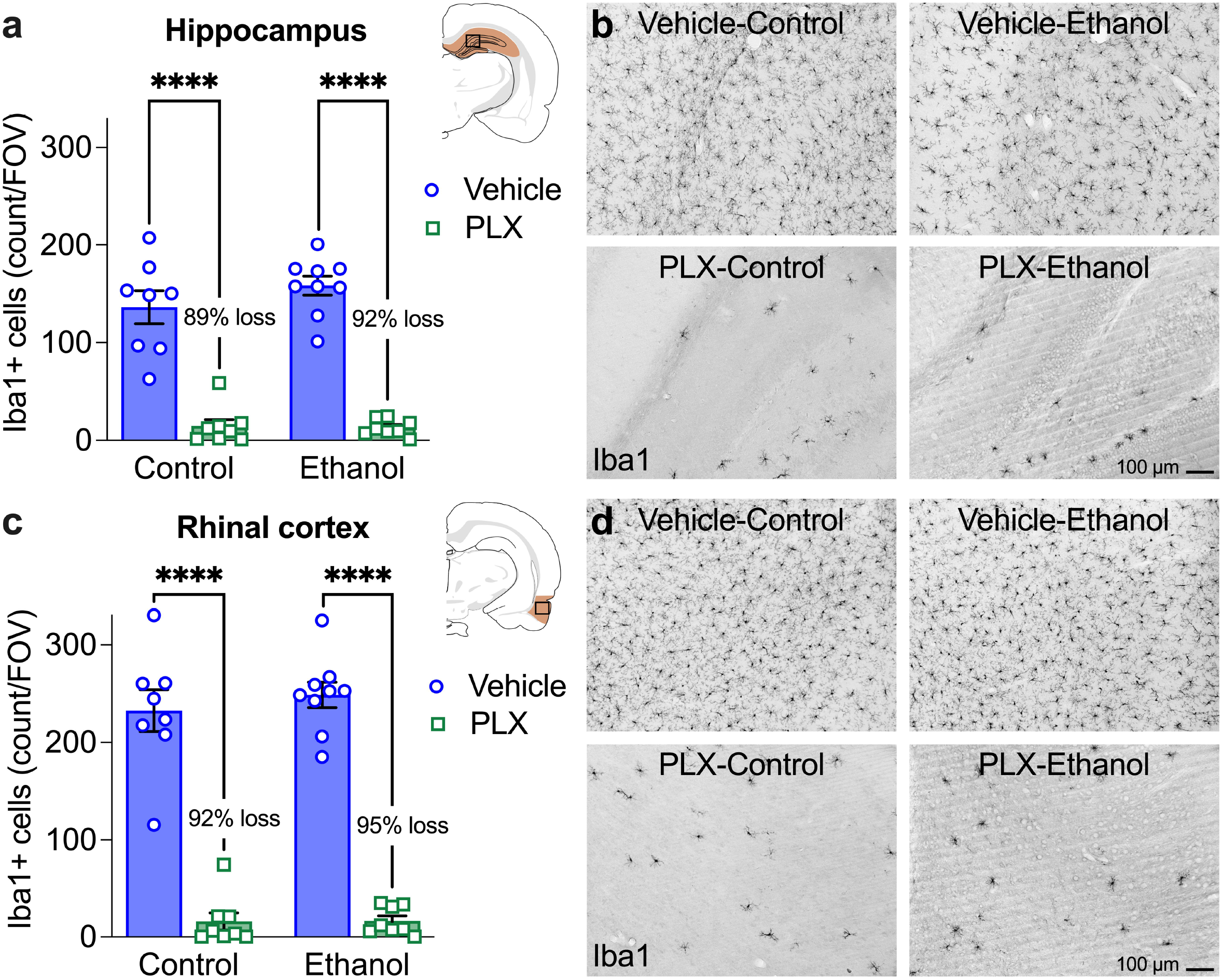
Robust microglia depletion with PLX5622 in adult male rats. PLX5622 administration reduced microglia counts (Iba1+) compared to vehicle in both diet control and ethanol groups in the hippocampus (**a**) and rhinal cortex (**c**). Representative images of microglia (Iba1+) in the hippocampus (**b**) and rhinal cortex (**d**) following treatment. FOV = field of view. \*\*\*\**p* < 0.0001

Microglia depletion with PLX5622 blocked ethanol-induced neurodegeneration, as the ethanol-induced increase in FJB+ (a marker of dying neurons) cells was reduced by PLX treatment (Figure 2). In the hippocampus, two-way ANOVA revealed main effects of drug (F_1,29_ = 4.883, *p* = 0.035) and diet (F_1,29_ = 5.777, *p* = 0.023) and an interaction (F_1,29_ = 5.699, *p* = 0.024). Posthoc comparisons show increased FJB+ cells in the ethanol-vehicle group compared to both the control-vehicle (*p* = 0.007) and ethanol-PLX (*p* = 0.010) groups. In the rhinal cortex, there were main effects of drug (F_1,29_ = 6.935, *p* = 0.013) and diet (F_1,29_ = 13.511, *p* = 0.001) and an interaction (F_1,29_ = 8.884, *p* = 0.006). Posthoc comparisons show FJB+ cells were increased in the ethanol-vehicle group compared to both the control-vehicle (*p* = 0.0002) and ethanol-PLX (*p* = 0.001) groups.

**Fig. 2.**
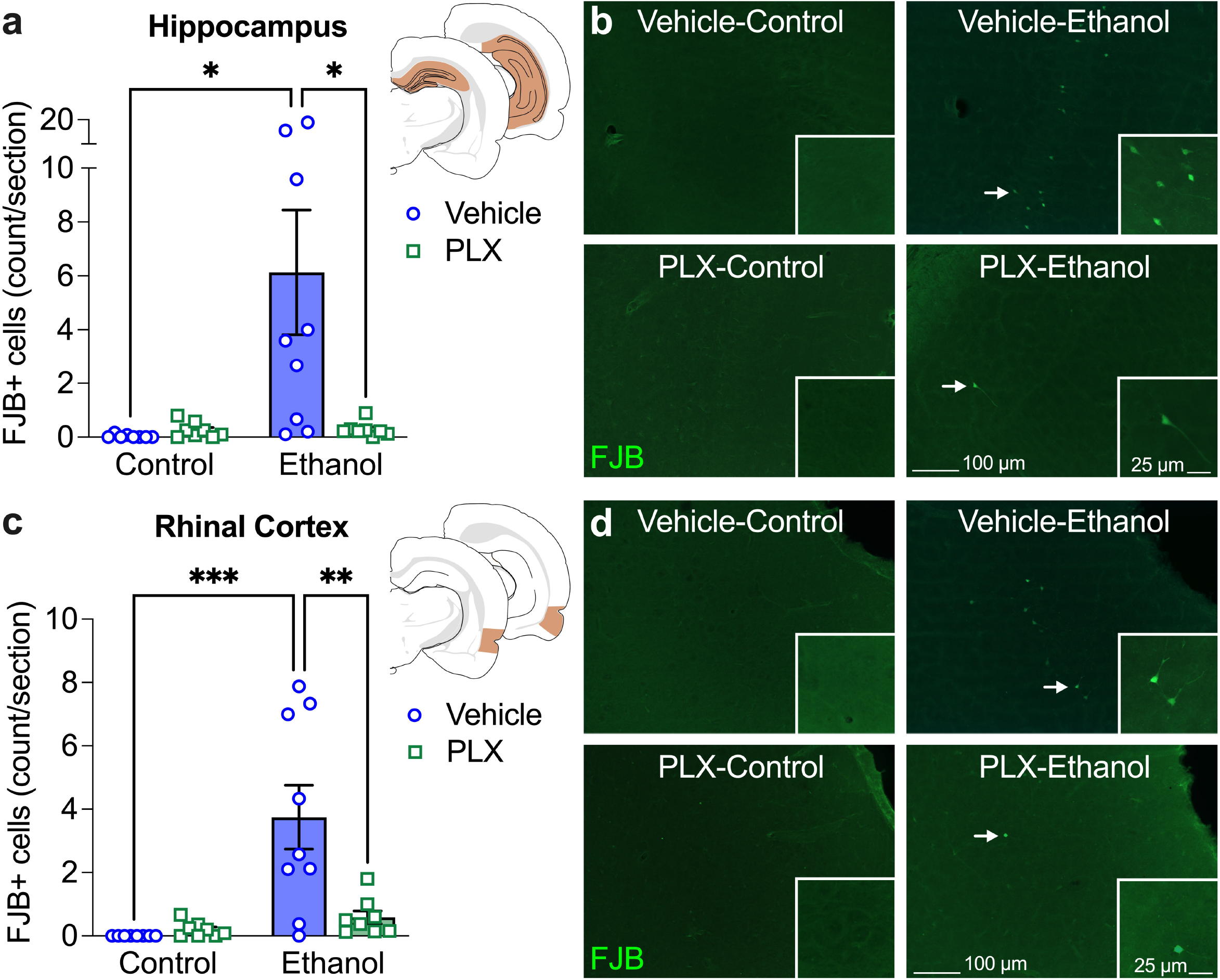
Lack of ethanol-induced neurodegeneration (FJB+) with PLX5622 in adult male rats. Binge-like ethanol exposure increased FJB+ cells in the hippocampus (**a**) and rhinal cortex (**c**) of vehicle-treated rats compared to the control-vehicle group and, critically, the ethanol-PLX group. Representative images of FJB staining (arrows) in the hippocampus (**b**) and rhinal cortex (**d**). \**p* < 0.05, \*\**p* < 0.01, \*\*\**p* < 0.001

Importantly, there was no effect of PLX5622 on intoxication behavior (ethanol-vehicle 1.8 ± 0.1; ethanol-PLX 1.9 ± 0.2; 0-5 scale; *p* = 0.774), ethanol dose (ethanol-vehicle 9.5 ± 0.4 g/kg/day; ethanol-PLX 9.3 ± 0.6 g/kg/day; *p* = 0.774), or BEC (ethanol-vehicle 399.1 ± 16.9 mg/dl; ethanol-PLX 381.9 ± 28.1 mg/dl; *p* = 0.609). Thus, PLX did not appear to impact ethanol pharmacokinetics.

### Discussion

This study is the first to directly implicate microglia in ethanol-induced neuronal death (FJB+) as PLX-treated rats exposed to ethanol showed minimal FJB+ cells versus ethanol-vehicle rats. This report is also the first to show full (>90%) microglia depletion with PLX5622 in a rat model of alcohol exposure, verified by immunohistochemistry for the pan-microglia marker, Iba1. Importantly, PLX5622 did not appear to impact ethanol pharmacokinetics, as BECs were similar between PLX- and vehicle-treated rats.

These findings provide direct evidence that microglia have a causal role in alcohol-induced neurodegeneration. A hyperactive neuroimmune response has long been implicated in alcohol-induced neurodegeneration, evidenced by increased proinflammatory cytokine expression in mouse models of AUD (reviewed in Crews and Vetreno, 2014; Erickson et al., 2019) and the role inflammatory cytokines play in alcohol-induced damage (Qin and Crews, 2012). Ties to microglia were supported by observations of increased expression of microglia markers and reactive morphology in postmortem brains of men with AUD (He and Crews, 2008) and similar findings across animal models (Marshall et al., 2013; Peng et al., 2017; Lowe et al., 2020; reviewed in Melbourne et al., 2019). Depleting microglia with PLX5622 blunted pro-inflammatory cytokine signaling following acute alcohol exposure in C57Bl6J mice (Walter and Crews, 2017), while depletion and repopulation with PLX3397, a less specific CSF1R antagonist, reduced most ethanol-induced neuroimmune gene expression in rat organotypic hippocampal slice culture (Coleman et al., 2020). However, neither study examined neurodegeneration.

The role of microglia in the neuroimmune hypothesis of AUD has been based primarily on findings in mice, as more subtle responses to alcohol occur in rats (reviewed in Melbourne et al., 2019). For example, only mild changes in microglia phenotype have been observed following alcohol exposure (Peng et al., 2017) with little evidence of cytotoxic reactivity according to histological, or protein or gene expression of microglia-specific markers, growth factors and cytokines (e.g. Marshall et al., 2013; reviewed in Melbourne et al., 2019). As such, these findings were somewhat unexpected in a rat model, but perhaps even subtle microglia responses are deleterious in the short term. These data suggest that during intoxication, microglia reactions have neurotoxic consequences, consistent with the aforementioned pro-inflammatory effects of microglia in some alcohol models (Qin and Crews, 2012; Walter and Crews, 2017; Melbourne et al., 2019; Coleman et al., 2020). Alternatively, the protective effects of depletion may be due, in part, to depletion of peripheral CSF1R-expressing cells, as peripheral macrophages may drive alcohol-induced neuroinflammatory events in mice (Lowe et al., 2020). However, there is little evidence that the blood-brain-barrier is damaged in the current rat model of an AUD (Marshall et al., 2013).

Microglia depletion strategies allow dissection of microglial-mediated mechanisms and have been used in several neurodegenerative disease models among others (Green et al., 2020). CSF1R antagonists have been used in mouse models of ethanol exposure and revealed important roles of microglia in the development of AUD. Specifically, microglia depletion prevented consequences of excessive ethanol consumption such as the acute withdrawal-induced increases in proinflammatory cytokine TNFα (Walter and Crews, 2017) but more critically, prevented the escalation of alcohol drinking, a hallmark of the development of an AUD (Warden et al., 2020).

As exciting as this discovery is for the field, there are some limitations though a host of future directions. First, these studies were conducted in males only, as initial testing in females indicated a potential interaction between ethanol and the vehicle/route of administration used for microglia depletion, requiring modification of delivery in future work. It is also important to consider astrocyte involvement in these processes, as prior work indicates microglia depletion prevents astrocyte response to insult (Riquier and Sollars, 2020). Lastly, the signaling mechanism of ethanol-microglia-induced neurodegeneration remains unknown, but there are several possible contenders. Alcohol induces microglia reactivity by activating toll-like receptors (TLRs), specifically TLR4, on microglia, and triggers release of danger-associated molecular patterns (DAMPs) following neuronal damage (Crews and Vetreno, 2014; Erickson et al., 2019). Several molecules implicated in ethanol-induced neurodegeneration are released by microglia, including TNF-α, IL-1β, IL-6, TGF-β1, and reactive oxidative species (ROS). Further research is needed to determine which molecule(s) mediate cell death in this model. Taken together, these findings make an exciting advance for the study of neuroprotection in AUD.

## Author contributions

Erika R. Carlson (E.R.C.) and Jennifer K. Melbourne (J.K.M.) have contributed equally to this research. J.K.M. and Kimberly Nixon (K.N.) conceptualized and designed the project; J.K.M. and E.R.C. performed method development, formal analyses, and data curation. K.N. and J.K.M. acquired funding. All authors contributed to data collection, interpretation of data, as well as writing, review, and editing the manuscript. All authors have reviewed and approve of the final manuscript.

## Statements and declarations

### Ethics statement

All animal use procedures were approved by The University of Texas at Austin Institutional Animal Care and Use Committee (IACUC) in advance and performed according to National Institutes of Health guidelines.

## References

Coleman LG, Jr., Zou J, Crews FT (2020) Microglial depletion and repopulation in brain slice culture normalizes sensitized proinflammatory signaling. J Neuroinflammation 17:27. 10.1186/s12974-019-1678-y

Crews FT, Vetreno RP (2014) Neuroimmune basis of alcoholic brain damage. Int Rev Neurobiol 118:315–357. 10.1016/B978-0-12-801284-0.00010-5

Erickson EK, Grantham EK, Warden AS, Harris RA (2019) Neuroimmune signaling in alcohol use disorder. Pharm Biochem Behav 177:34–60. 10.1016/j.pbb.2018.12.007

Green KN, Crapser JD, Hohsfield LA (2020) To Kill a Microglia: A Case for CSF1R Inhibitors. Trends Immunol 41:771–784. 10.1016/j.it.2020.07.001

He J, Crews FT (2008) Increased MCP-1 and microglia in various regions of the human alcoholic brain. Exp Neurol 210:349–358. 10.1016/j.expneurol.2007.11.017

Kelso ML, Liput DJ, Eaves DW, Nixon K (2011) Upregulated vimentin suggests new areas of neurodegeneration in a model of an alcohol use disorder. Neuroscience 197:381–393. 10.1016/j.neuroscience.2011.09.019

Lowe PP, Morel C, Ambade A, Iracheta-Vellve A, Kwiatkowski E, Satishchandran A, Furi I, Cho Y, Gyongyosi B, Catalano D, Lefebvre E, Fischer L, Seyedkazemi S, Schafer DP, Szabo G (2020) Chronic alcohol-induced neuroinflammation involves CCR2/5-dependent peripheral macrophage infiltration and microglia alterations. J Neuroinflammation 17:296. 10.1186/s12974-020-01972-5

Marshall SA, McClain JA, Kelso ML, Hopkins DM, Pauly JR, Nixon K (2013) Microglial activation is not equivalent to neuroinflammation in alcohol-induced neurodegeneration: The importance of microglia phenotype. Neurobiol Dis 54:239–251. 10.1016/j.nbd.2012.12.016

Melbourne JK, Thompson KR, Peng H, Nixon K (2019) Its complicated: The relationship between alcohol and microglia in the search for novel pharmacotherapeutic targets for alcohol use disorders. In: Molecular Basis of Neuropsychiatric Disorders: from Bench to Bedside, 1 Edition, pp 1–43: Elsevier Inc. 10.1016/bs.pmbts.2019.06.011

Peng H, Geil Nickell CR, Chen KY, McClain JA, Nixon K (2017) Increased expression of M1 and M2 phenotypic markers in isolated microglia after four-day binge alcohol exposure in male rats. Alcohol 62:29–40. 10.1016/j.alcohol.2017.02.175

Qin L, Crews FT (2012) NADPH oxidase and reactive oxygen species contribute to alcohol-induced microglial activation and neurodegeneration. J Neuroinflammation 9:5. 10.1186/1742-2094-9-5

Riquier AJ, Sollars SI (2020) Astrocytic response to neural injury is larger during development than in adulthood and is not predicated upon the presence of microglia. Brain Behav Immun Health 1:100010. 10.1016/j.bbih.2019.100010

Walter TJ, Crews FT (2017) Microglial depletion alters the brain neuroimmune response to acute binge ethanol withdrawal. J Neuroinflammation 14:86. 10.1186/s12974-017-0856-z

Warden AS, Wolfe SA, Khom S, Varodayan FP, Patel RR, Steinman MQ, Bajo M, Montgomery SE, Vlkolinsky R, Nadav T, Polis I, Roberts AJ, Mayfield RD, Harris RA, Roberto M (2020) Microglia Control Escalation of Drinking in Alcohol-Dependent Mice: Genomic and Synaptic Drivers. Biological psychiatry 88:910–921. 10.1016/j.biopsych.2020.05.011

Zahr NM, Pfefferbaum A (2017) Alcohol’s Effects on the Brain: Neuroimaging Results in Humans and Animal Models. Alcohol Research: Current Reviews 38:183–206.

